# VPS41 loss triggers iron overload, oxidative stress, and mitochondrial fragmentation linked to ferroptosis

**DOI:** 10.64898/2026.05.15.725396

**Authors:** Reini E. N. van der Welle, Rebecca Jark, Judith J.M. Jans, Nanda M. Verhoeven-Duif, Judith Klumperman

## Abstract

The tight regulation of iron homeostasis is of great importance for cellular health. An increase in intracellular iron levels results in the formation of free radicals, which damages macromolecules and membranes, eventually resulting in cell death by Ferroptosis. Recently, we showed that patients with mutations in VPS41 display a severe neurodegenerative phenotype with iron deposition in the brain. VPS41 is well known as subunit of the HOPS complex required for fusion of late endosomes and autophagosomes with lysosomes. However, VPS41 has also been identified as inhibitor of Ferroptosis and regulator of redox homeostasis. How VPS41 exerts these functions and if these are dependent on the HOPS complex is unknown. Here we show that depletion of VPS41 results in increased intracellular iron levels, ROS formation and mitochondrial fission. Our findings indicate an important role for VPS41 in the regulation of iron homeostasis and mitochondrial fission and suggest Ferroptosis as a possible cause for neurodegeneration in VPS41 patients.

## Introduction

Iron is a vital nutrient for a variety of cellular functions, such as lipid metabolism, oxygen transport, DNA synthesis and respiration. Since iron is toxic at high concentrations, a tight control of iron homeostasis is required(Mackenzie et al., 2008; Sousa et al., 2020). Iron enters the cell as ferric iron (Fe^3+^) bound to transferrin (Tf), which is taken up by the transferrin receptor (TfR). Upon internalization, the slightly lower pH in early endosomes causes dissociation of iron from Tf, upon which ferric Fe^3+^ is converted into ferrous Fe^2+^, the highly reactive configuration of iron(C. P. Anderson et al., 2012; G. J. Anderson & Frazer, 2017; Mayle et al., 2012). Via the Divalent Metal Transporter 1 (DMT1), Fe^2+^ is transported across the endosomal membrane into the cytosol. Here it can be converted to Fe^3+^ and stored in complex with Ferritin, the major iron storage protein(Kakhlon & Cabantchik, 2002; Kidane et al., 2006). Alternatively, cytosolic Fe^2+^ can be transported into mitochondria to form iron-sulfur clusters (Fe-S) required for electron transfer within the mitochondrial respiratory chain(Read et al., 2021; Stehling & Lill, 2013). Lastly, cytosolic Fe^2+^ can form a labile iron pool. This pool is ready for instant use but becomes toxic when Fe^2+^ levels rise too high(Kakhlon & Cabantchik, 2002; Lv & Shang, 2018). To regulate this, Fe^2+^ can be released from cells via Ferroportin, the only known iron exporter present in the cell membrane. Upon release into the extracellular environment, the multicopper oxidase Ceruloplasmin converts Fe^2+^ to Fe^3+^ to enable binding to Tf again(De Domenico et al., 2007; Donovan et al., 2005; Musci et al., 2014).

The expression levels of proteins involved in iron metabolism is controlled by iron itself, via the Iron Regulatory Proteins 1 and 2 (IRP1 and IRP2). These IRPs bind Iron Responsive Elements (IREs) present in the mRNAs of proteins involved in iron uptake (Tf and TfR, DMT1), export (Ferroportin) and storage (Ferritin). Interestingly, IRP1 and IRP2 sense the cytosolic iron concentration and modify expression of their target mRNAs correspondingly. IRP binding to mRNAs encoding Ferritin and Ferroportin results in their repressed translation and increased degradation. Contrarily, IRP stabilizes the mRNAs encoding for TfR and DMT1, thereby inhibiting degradation and promoting translation. Low iron conditions induce IRP-IRE binding, upregulating TfR and DMT1 translation to allow increased iron uptake from the extracellular environment and endosomal export, respectively. By contrast, translation of Ferritin and Ferroportin is repressed, ensuring reduced iron storage and cellular iron export. In case of iron overload, IRP-IRE binding is inhibited, resulting in increased Ferritin and Ferroportin translation, but degradation of TfR and DMT1. Thus, the IRP/IRE system is key for maintaining cellular iron homeostasis(C. P. Anderson et al., 2012; Wilkinson & Pantopoulos, 2014) and understanding its regulation is of great importance in light of the many iron-associated pathologies that derive from iron deficiency or overload (See also Chapter 1, Figure 2 and 3).

**Figure 1.**
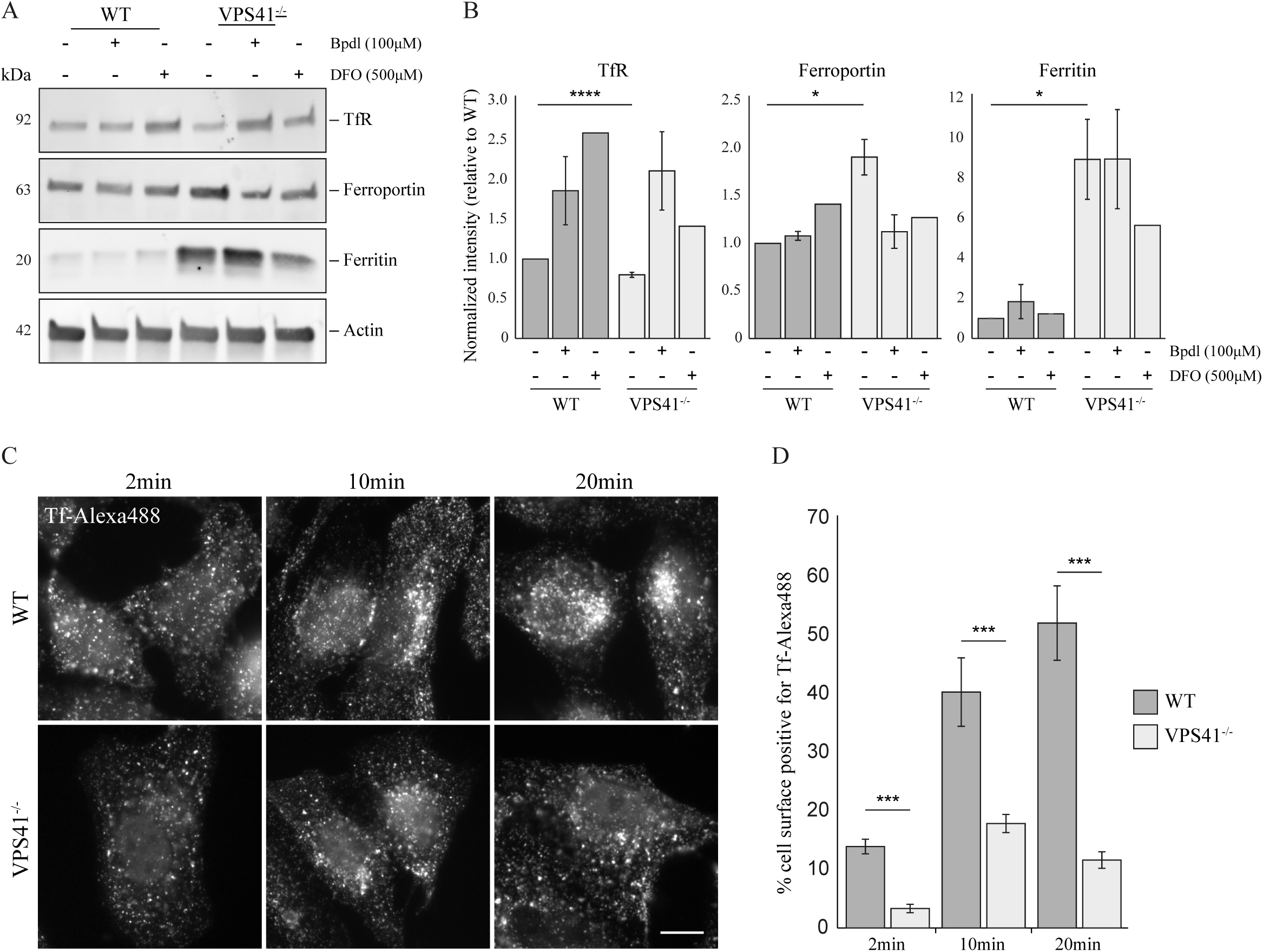
Iron-associated proteins all indicate increased iron levels in VPS41^-/-^ cells. A. Western blot on Transferrin Receptor (TfR), Ferroportin and Ferritin protein levels under normal and iron chelated (Bpdl and DFO) conditions. TfR protein levels are decreased, whereas Ferroportin and Ferritin levels are increased in VPS41-/- cells under basal conditions. Upon iron chelation, TfR levels are increased while Ferroportin and Ferritin levels are reduced, indicative of functional IRE/IRP regulation of these iron-associated proteins in VPS41-/- cells. B. Quantification of A, bars represent mean ± SEM, *P < 0.05, ****P < 10^-5^. One-way ANOVA with Bonferroni correction) (n=2). C. Transferrin (Tf) uptake in WT versus VPS41^-/-^ cells, assessed after 2, 10 and 20 minutes Tf-Alexa488 incubation. VPS41^-/-^ cells show reduced Tf uptake for all time points compared to WT cells. (Scale bare; 10µm) D. Quantification of C, showing percentage of cell surface positive for Tf-Alexa488 signal. Bars represent mean ± SEM, ***P < 0.001. One-way ANOVA with Bonferroni correction) (n=3).

**Figure 2.**
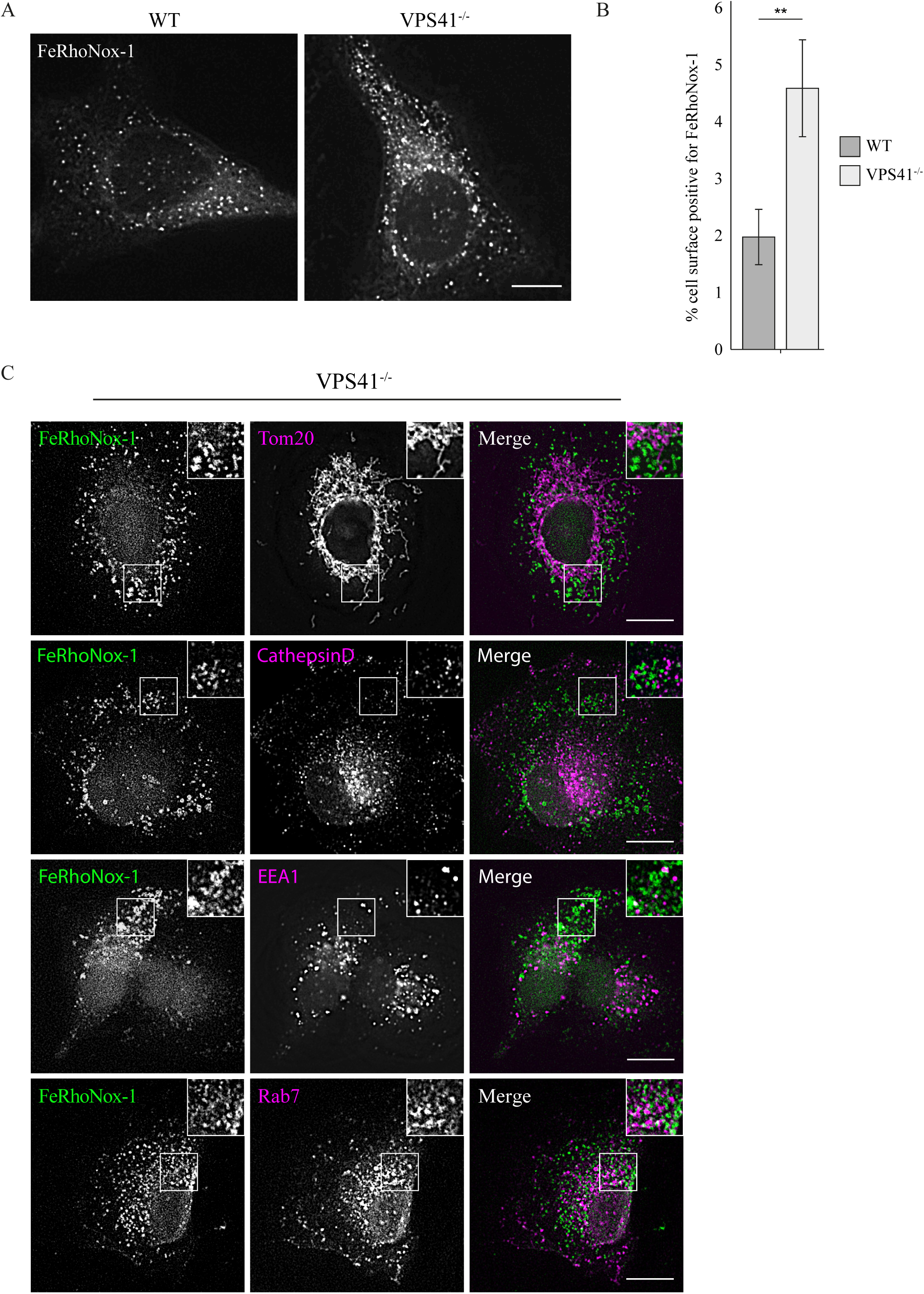
The labile iron pool is increased in VPS41^-/-^ cells. A. Visualization of ferrous (Fe^2+^) iron using the FeRhoNox-1 probe shows an increase in labile iron in VPS41^-/-^ cells compared to WT cells. (Scale bar; 10µm). B. Quantification of A, showing percentage of cell surface positive for FeRhoNox-1 signal. Bars represent mean ± SEM, **P < 0.01, Unpaired t-test, n=2). C. Immunofluorescence of endogenous EEA1, Rab7, Cathepsin D and Tomm20 in VPS41^-/-^ cells incubated with FeRhoNox-1 to visualize localization of Fe^2+^ clusters. No co-localization was observed between FeRhoNox-1 and Tomm20 or Cathepsin D, whereas some co-localization of FeRhoNox-1, EEA1 and Rab7 was found. (Scale bar; 10µm) (n=1).

**Figure 3.**
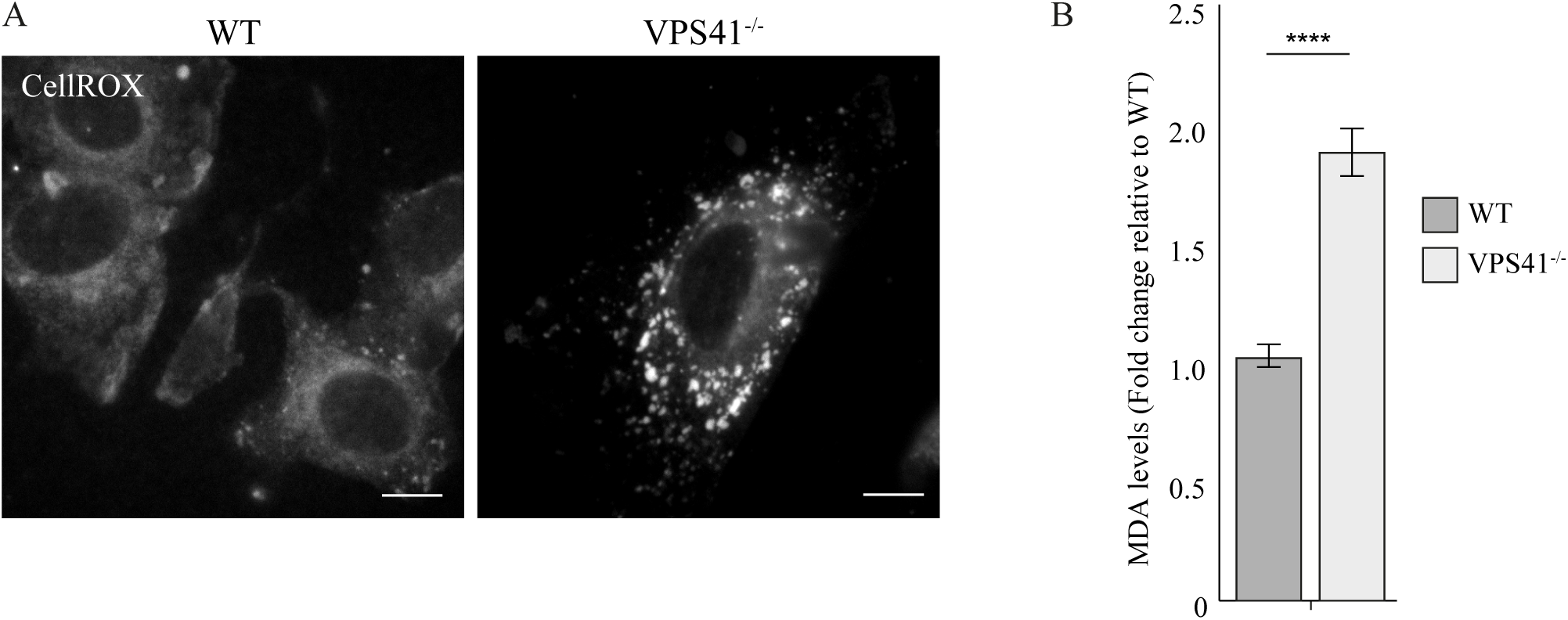
VPS41^-/-^ cells show increased oxidative stress levels. A. Visualization of Reactive Oxygen Species (ROS) in WT and VPS41^-/-^ cells using the CellROX™ Deep Red probe. WT cells show no distinct signal, whereas cell depleted of VPS41 display ROS positive clusters. B. Lipid peroxidation was assessed by measuring the malondialdehyde (MDA) levels in WT and VPS41^-/-^cells. MDA levels were significantly increased upon deletion of VPS41. Bars represent mean ± SEM, ****P < 10^-5^, Unpaired t-test, n=2).

When cellular iron levels are high, the redox homeostasis - the balance between oxidation and reduction reactions - is disrupted, which leads to the production of Reactive Oxygen Species (ROS)(Nakamura et al., 2019; Ying et al., 2021). These free radicals are generated by the Fenton chain reaction, which encompasses a series of reactions between peroxides and Fe^2+^. A consequence of the Fenton reaction and ROS production is the peroxidation of phospholipids containing polyunsaturated fatty acids(Rynkowska et al., 2020; Sun et al., 2018). This compromises membranes and damages macromolecules, eventually resulting in Ferroptosis, a relatively recent discovered type of regulated cell death(X. Jiang et al., 2021). Ferroptosis is controlled by the same regulatory mechanisms as involved in iron homeostasis and is associated with various iron-related pathologies, such as cancer, ischemia-reperfusion and neurodegenerative diseases(Guiney et al., 2017; Ji et al., 2022). Intriguingly, a recent study identified depletion of VPS41 as initiator of Ferroptosis(Tian et al., 2021). VPS41 is best known for its role in the Homotypic fusion and protein sorting (HOPS) complex, a multisubunit tethering complex required for lysosomal fusion events. However, VPS41 was originally described in yeast as protein necessary for growth on low-iron medium, indicating a role in high-affinity iron uptake(Balderhaar & Ungermann, 2013; Lu et al., 2007; Radisky et al., 1997; Solinger & Spang, 2013). Moreover, MRI measurements by us and others recently showed that compound heterozygote mutations in VPS41 cause a neurodegenerative disease with iron accumulation in the basal ganglia, globus pallidus or substantia nigra(Sanderson et al., 2021; Steel et al., 2020; Van der Welle et al., 2021).

As part of the HOPS complex, VPS41 is required for homo- and heterotypic fusion events between lysosomes and late endosomes or autophagosomes(Balderhaar & Ungermann, 2013; J. Van Der Beek et al., 2019; P. Jiang et al., 2014; McEwan et al., 2015; Pols, Ten Brink, et al., 2013; Shvarev et al., 2022). In addition, VPS41 can recruit TGN-derived vesicles carrying lysosomal membrane proteins to late endosomes. Subsequent HOPS-dependent fusion establishes a pathway to selectively modulate the molecular composition of late-stage endosomal compartments(Sanzà et al., 2025). Furthermore, HOPS is involved in the regulation of ‘mammalian target of rapamycin complex 1’ (mTORC1) signaling(Van der Welle et al., 2021). The mTORC1 kinase complex is a master regulator of cellular growth, of which the activity is controlled by extra- and intracellular stimuli, like nutrients, growth factors and energy(Rabanal-Ruiz & Korolchuk, 2018; Zoncu et al., 2011). Emerging evidence shows the existence of a canonical and non-canonical mTORC1 signaling pathway, enabling a selective response towards mTORC1 substrates controlling growth or metablolism(Napolitano et al., 2020, 2022). Previously, we showed that mutations in HOPS subunits specifically impair the non-canonical mTORC1 pathway, which regulates autophagy and lysosomal biogenesis(Van der Welle et al., 2021). Recently, we found that HOPS is required to recruit the Rag/Folliculin (FLCN) complex to endo-lysosomes, which is necessary for phosphorylation of the MiTF/TFE transcription factors TFE3 and TFEB (van der Welle et al., Chapter 3, manuscript under preparation) (Martina & Puertollano, 2013; Napolitano et al., 2020, 2022; Puertollano et al., 2018). In the absence of HOPS, non-phosphorylated TFE3/TFEB translocate to the nucleus, where they promote expression of the Coordinated Lysosomal Expression and Regulation (CLEAR) gene network, inducing autophagy and lysosome biogenesis(Palmieri et al., 2011; Roczniak-ferguson et al., 2012). Interestingly, various proteins involved in mTORC1 signaling, e.g. FLCN, AMP-activated protein kinase (AMPK) and TFEB, are also implicated in iron homeostasis and ROS production. FLCN is identified as interactor of Rab11A and is therefore involved in TfR recycling, whereas chelation of iron inhibits canonical mTORC1 signaling by activating the AMPK pathway(Fernández et al., 2022; Ramirez Reyes et al., 2021; Shang et al., 2020; Wang et al., 2021; Zhang et al., 2016).

In this chapter, we aimed to further unravel the role of mammalian VPS41 in iron homeostasis and oxidative stress, by investigating the effects of VPS41 depletion on intracellular iron pools in human cells. We show that depletion of VPS41 reduces TfR and elevates Ferritin and Ferroportin protein levels, as well as the labile iron pool. These are all indicators for increased intracellular iron levels. Additionally, we found that VPS41^-/-^ cells showed an increase in ROS levels and a concomitant increase in Drp1-mediated mitochondrial fission. Interestingly, these effects were specific for VPS41 and not found after depletion of other HOPS members. Our study thus highlights a specific role for VPS41 in redox signaling and iron homeostasis with potential implications for the onset of Ferroptosis.

## Results

### VPS41^-/-^ cells show increased intracellular iron levels

To address the role of VPS41 in iron metabolism, we made A549 cells knocked out for VPS41 by CRISPR/Cas (VPS41^-/-^). To assess the iron load in these cells, we analyzed the intracellular levels of Transferrin Receptor (TfR), Transferrin (Tf), Ferritin and Ferroportin. These iron-related proteins are generally used as indicator for iron load, since direct analysis of intracellular iron concentrations has been proven difficult(C. P. Anderson et al., 2012; Mackenzie et al., 2008).

By Western blot analysis we found that TfR levels were significantly decreased in VPS41^-/-^ compared to wildtype (WT) cells (Fig 1A). Accordingly, when WT and VPS41^-/-^ cells were incubated with Tf-Alexa488 for 2, 10 or 20 minutes, VPS41^-/-^ cells contained lower Tf levels at all timepoints (Fig 1B). Since TfR and Tf are required for iron uptake, both results were indicative for a lowering of the uptake of iron by VPS41^-/-^ cells, which is a typical response to iron overload. Of note, in a previous study where we knocked down (KD) VPS41 using RNA interference (RNAi), we found no effect on Tf uptake(Pols, Ten Brink, et al., 2013). This discrepancy can be attributed to the long-term depletion of VPS41 in VPS41^-/-^ cells compared to only 3 days of KD conditions.

Ferritin together with the iron exporter Ferroportin control the labile iron pool of reactive Fe^2+^ (Kakhlon & Cabantchik, 2002; Kidane et al., 2006; Lv & Shang, 2018). Western blot analysis of these proteins showed that VPS41^-/-^ cells contained significant higher Ferritin and Ferroportin protein levels (Fig 1A). Since increased Ferritin and Ferroportin expression leads to a reduction of the labile iron pool, these data were again indicative for iron overload in VPS41^-/-^ cells.

To assess if VPS41^-/-^ cells were still able to respond to changes in intracellular iron levels, we treated them with 100 μM 2,2’-bipyridyl (Bpdl) or 500 μM Deferoxamine (DFO), two iron chelators capable of reducing iron overload. Although the effectiveness between the Bpdl and DFO treatment varied, in general iron chelation increased TfR and decreased Ferroportin and Ferritin protein levels in VPS41^-/-^cells (Fig 1B). Thus, VPS41^-/-^ cells were capable to respond to a change in iron levels, indicating that regulation of iron homeostasis by the IRP/IRE system was not affected.

Together these data indicated that VPS41 depletion led to increased intracellular iron levels, to which cells responded by altering the expression levels of iron controlling proteins.

### Depletion of VPS41 increases the labile iron pool

A key feature of Ferroptosis is an increased redox-prone, labile iron pool, which mainly consists of ferrous (Fe^2+^) iron(Kakhlon & Cabantchik, 2002; Lv & Shang, 2018). The assays on iron-associated proteins (Fig. 1A and B) indicated an increase in intracellular iron but gave no specific information on the labile iron pool. To address this, WT and VPS41^-/-^ cells were incubated with FeRhoNox-1, a probe that only becomes fluorescent upon binding to Fe^2+^. Of note, FeRhoNox-1 is membrane permeable and binds the cytosolic labile iron pool as well as Fe^2+^ present in endosomes(Halcrow et al., 2022). In WT cells, the FeRhoNox-1signal appeared as a dispersed, punctate fluorescence pattern. Interestingly, in VPS41^-/-^ cells this signal was significantly enhanced (Fig 2A, B). To identify the nature of the FeRhoNox-1-positive spots, we co-labeled cells for the mitochondrial marker Tomm20 or the endo-lysosomal marker Cathepsin D. Neither of these antibodies showed noticeable overlap with the FeRhoNox-1 spots (Fig 2C). Next, we labeled for the early endosomal marker EEA1 and late endosomal marker Rab7. In WT cells these markers localize to distinct populations of endosomes (Fig 2C), whereas in VPS41^-/-^ cells they re-localize to enlarged, hybrid early-late endosomal compartments that are induced by depletion of the HOPS complex (further referred to as HOPS bodies)(van der J. Beek et al., 2022). In WT cells, FeRhoNox-1 only mildly co-localized with either EEA1 or Rab7, but inVPS41^-/-^ cells FeRhoNox-1 partially co-localized with EEA1 and Rab7 in the HOPS bodies. Most likely, the FeRhoNox-1 spots that did not co-localize with any of the marker proteins represented the cytosolic pool of Fe^2+^.

We concluded from these experiments that VPS41^-/-^ depletion induced increased levels of the labile iron pool of reactive Fe^2+^ in the cytosol as well as within aberrant early-late endosomes induced by HOPS depletion.

### VPS41^-/-^ and VPS41 patient cells show increased oxidative stress levels

An increase in Fe^2+^ levels is commonly associated with oxidative stress, since Fe^2+^ catalyzes the formation of toxic reactive oxygen species (ROS) via the Fenton reaction(Nakamura et al., 2019; Rynkowska et al., 2020). To determine if VPS41 depletion induced ROS, WT and VPS41^-/-^ cells were incubated with cellROX Deep Red (Fig 3A). This probe is non-fluorescent in its reduced state but emits a fluorescent signal upon oxidation by ROS. WT cells treated with cellROX showed a weak, cytosolic signal. Strikingly, in VPS41^-/-^ cells this signal was manifold stronger (Fig 3B) and, interestingly, appeared as clusters rather than dispersed (Fig 3A). Additional experiments are required to determine if these clusters represented cytosolic aggregates or organelles.

A major implication of increased ROS levels is lipid peroxidation, a key driver of Ferroptosis(Feng & Stockwell, 2018). Malondialdehyde (MDA), one of the final products of peroxidated polyunsaturated fatty acids, is often used as read-out for lipid peroxidation(Ayala et al., 2014). We assessed the MDA levels in WT and VPS41^-/-^ cells by quantifying MBA-TBA adduct levels, using a colorimetrical approach on cell lysates incubated with thiobarbituric acid (TBA). Interestingly, MDA levels were doubled in VPS41^-/-^ cells compared to WT cells, indicating that VPS41 depletion indeed caused a highly significant increase in lipid peroxidation (Fig 3B).

An increase in lipid peroxidation is the net result of glutathione (GSH) depletion and/or Glutathione Peroxidase 4 (GPx4) inhibition, which together catalyze the reduction of lipid peroxides. GSH is the major scavenger of free radicals and a key regulator of cellular redox balance. Diminished GSH levels by oxidation to glutathione disulfide (GSSG) increases the vulnerability of cells towards oxidative stress(Desideri et al., 2012; Doll et al., 2017). To examine the effect of VPS41 depletion on GSH levels, we performed metabolomics on primary fibroblasts originating from a VPS41 patient (VPS41^S285P/R662*^) or a healthy, non-related, individual. Previously we showed that both VPS41^S285P^ and VPS41^R662*^ are unable to form the HOPS complex(Van der Welle et al., 2021). Interestingly, by untargeted metabolomics we found that GSH levels were strikingly reduced in VPS41 patient cells (data not shown). These preliminary data again indicated increased oxidative stress levels in the absence of functional VPS41.

Taken together, the cellROX, MDA and GSH data all indicated that cells depleted of VPS41 show elevated oxidative stress. This included increased lipid peroxidation, which is one of the main drivers of Ferroptosis.

### VPS41 depleted cells contain more fragmented but less mitochondria

Mitochondrial fragmentation is commonly seen in cells suffering from elevated oxidative stress. Mitochondria are key organelles of iron utilization as main site for Fe-S cluster synthesis and maintenance of redox homeostasis. Although it is not yet confirmed whether mitochondria play a role in the onset of Ferroptosis, mitochondrial morphological changes, e.g., fragmentation and cristae enlargement, have been described in Ferroptotic cells(M. Gao et al., 2019; Neitemeier et al., 2017; Stehling & Lill, 2013). Also, mitochondrial fragmentation and dysfunction are associated with cognitive impairment and neurodevelopmental disorders(Huang et al., 2018; Lezi & Swerdlow, 2012).

To investigate the effect of VPS41 depletion on mitochondria morphology, WT and VPS41^-/-^ A549 cells were incubated with MitoTracker and processed for fluorescence microscopy (Fig 4A). While WT cells contained an intricate network of elongated mitochondria, in VPS41^-/-^ cells, mitochondria were shorter, more fragmented, and cells lacked typical elongated mitochondria (Fig 4A). A similar phenotype was seen in WT versus VPS41^S285P/R662*^ fibroblasts (Fig 4C). Quantification of the fluorescent images confirmed a significant decrease in the average mitochondrial length in A549 VPS41^-/-^ cells (Fig 4B).

**Figure 4.**
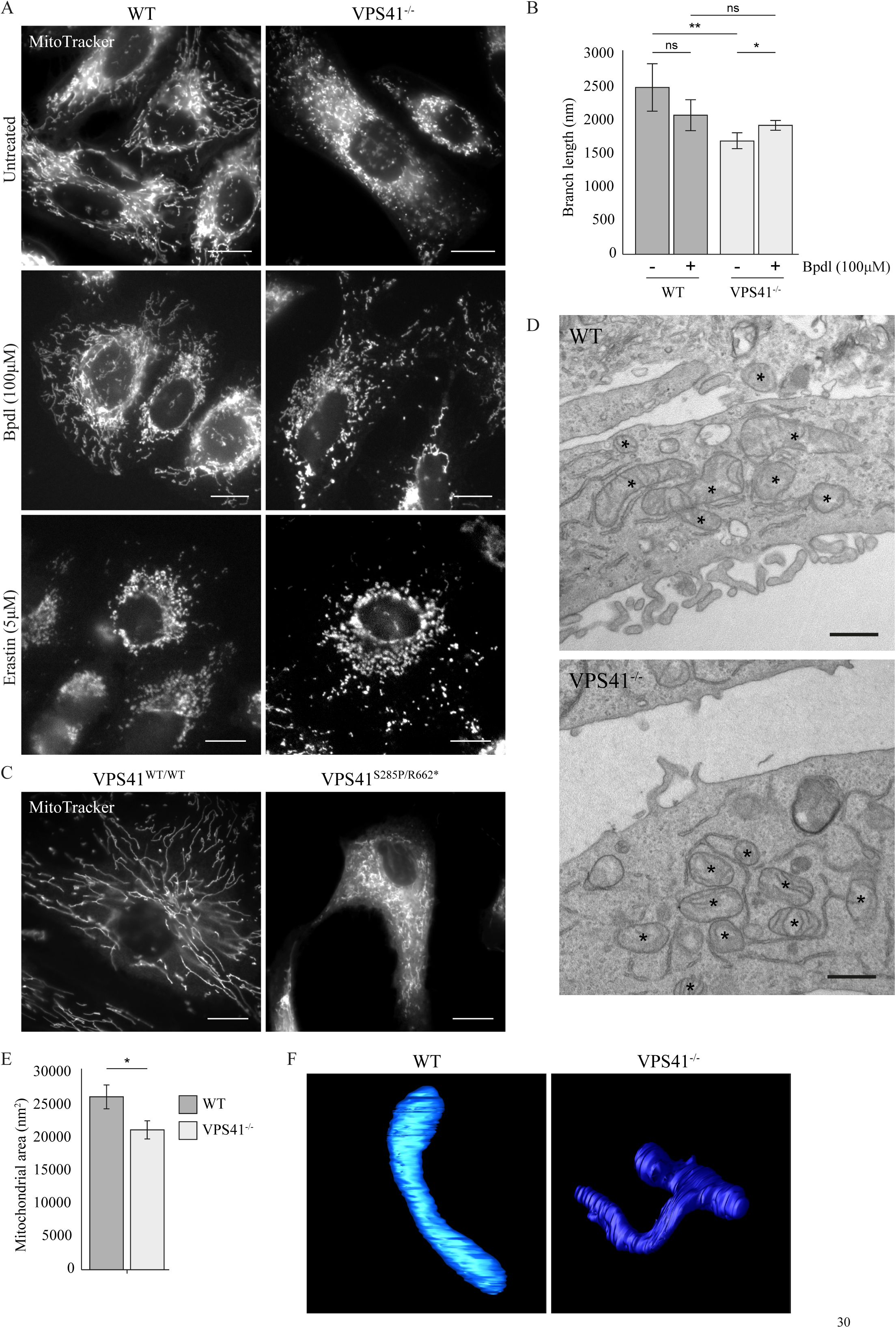
Mitochondrial morphology is affected in VPS41^-/-^ cells. A. Visualization of mitochondria in WT and VPS41^-/-^ cells using the MitoTracker probe under basal (Untreated), Ferroptosis induced (Erastin treatment) and iron chelated (Bpdl treatment) conditions. Under basal conditions, mitochondria in WT cells show elongated compartments, whereas VPS41^-/-^ cells lack these extended structures. Upon iron chelation, the fragmented mitochondria observed in VPS41^-/-^ is restored to some extent. Erastin treatment results in a fragmented mitochondrial phenotype in WT cells, comparable to VPS41^-/-^ cells. (Scale bar; 10µm). B. Quantification of the mitochondrial size under basal or iron chelated (Bpdl) conditions in WT and VPS41^-/-^ cells. Mitochondria are significantly smaller in VPS41^-/-^ cells compared to WT cells, which is restored upon Bpdl treatment. Bars represent mean ± SEM, *P < 0.05, **P < 0.01, One-way ANOVA with Bonferroni correction, n=2). C. Visualization of mitochondria in patient derived (VPS41^S285P/R662*^) and WT fibroblasts (VPS41^WT/WT^) using the MitoTracker probe. Mitochondria in patient cells are more fragmented and lack the elongated structure observed in WT fibroblasts. (Scale bar; 10µm, n=1). D. Representative electron micrographs of Epon sections of WT and VPS41^-/-^ cells. Smaller mitochondria are observed in VPS41^-/-^ cells, with absence of elongated structures. (Scale bar; 500nm). E. Quantification of mitochondrial area per cell surface on Epon sections of WT and VPS41^-/-^ cells. Mitochondria are smaller in cells depleted of VPS41. Bars represent mean ± SEM, *P < 0.05, Unpaired t-test, n=20 cells). F. 3D reconstruction by FIB-SEM of a mitochondrion of WT and VPS41^-/-^ cells. Mitochondria in VPS41^-/-^cells are smaller and less uniform in size with more branches compared to WT cells.

A similar mitochondrial phenotype has been described after oxidative stress, iron overload and Ferroptosis(Bauckman et al., 2013; M. Gao et al., 2019). To assess if iron chelation could rescue the alterations in mitochondrial morphology, WT and VPS41^-/-^ cells were treated with 100µM Bpdl (Fig 4A). This resulted in a significant increase in mitochondrial size in VPS41^-/-^ cells, whereas no significant change was observed for WT cells (Fig 4B). Interestingly, treatment of WT cells with Erastin, which induces Ferroptosis, resulted in a similar mitochondrial fragmentation phenotype as in VPS41^-/-^ cells (Fig 4A).

To assess the mitochondrial morphology by ultrastructural detail and to verify the phenotype in different cell lines, we prepared WT and VPS41^-/-^ HeLa cells for Electron Microscopy (EM) (Fig 4D). Notably, we found no obvious abnormalities in mitochondrial morphology, like aberrant cristae, density of the matrix or swelling, but morphometrical analysis showed that the total mitochondrial area per cell had decreased by ∼20% in VPS41^-/-^ cells (Fig 4E). This was a similar trend as in A459 cells and fibroblasts. Finally, to capture mitochondrial morphology in 3D, we performed Focused Ion Beam Scanning Electron Microscopy (FIB-SEM) of HeLa WT and VPS41^-/-^ mitochondria (Fig 4F and FIB-SEM dataset presented in (Fermie et al., 2018)). This revealed that mitochondria in VPS41^-/-^ were smaller in size, displayed a higher level or branching and were less uniform in shape.

Concluding, these data showed that depletion of VPS41 resulted in a fragmented mitochondrial network with a decrease in total mitochondrial volume. A similar phenotype was observed in WT cells after induction of Ferroptosis by Erastin. Additionally, the fragmented mitochondrial phenotype could be rescued in VPS41^-/-^ cells by iron chelation. Together these data indicated that depletion of VPS41 induced a strong mitochondrial phenotype as a result of iron overload.

### VPS41 depleted cells undergo increased mitochondrial fission

Small-sized mitochondria with increased branching and variable shapes commonly point to increased mitochondrial fission events(Westrate et al., 2014). Mitochondrial fission is coordinated by dynamin-related protein 1 (Drp1). Upon activation, this GTPase is recruited from the cytosol to the mitochondrial surface, where it polymerizes into a ring-like structure. When the GTPase domains of adjacent Drp1 molecules are in close proximity, hydrolysis takes place, causing polymer constriction and tightening of the Drp1 ring. Eventually, this results in fission of the mitochondria(F. Gao et al., 2021; Park et al., 2015a).

To determine if Drp1-dependent mitochondrial fission was increased in VPS41^-/-^ cells, we double-labeled A549 cells for endogenous Drp1 and MitoTracker (Fig 5A). In WT cells, Drp1 had a cytosolic distribution. By contrast, in VPS41^-/-^ cells Drp1 clearly co-localized with MitoTracker on mitochondria (Fig 5A, B). Interestingly, when we induced Ferroptosis in WT cells by Erastin, this also resulted in increased co-localization between Drp1 and MitoTracker (Fig 5A, B).

**Figure 5.**
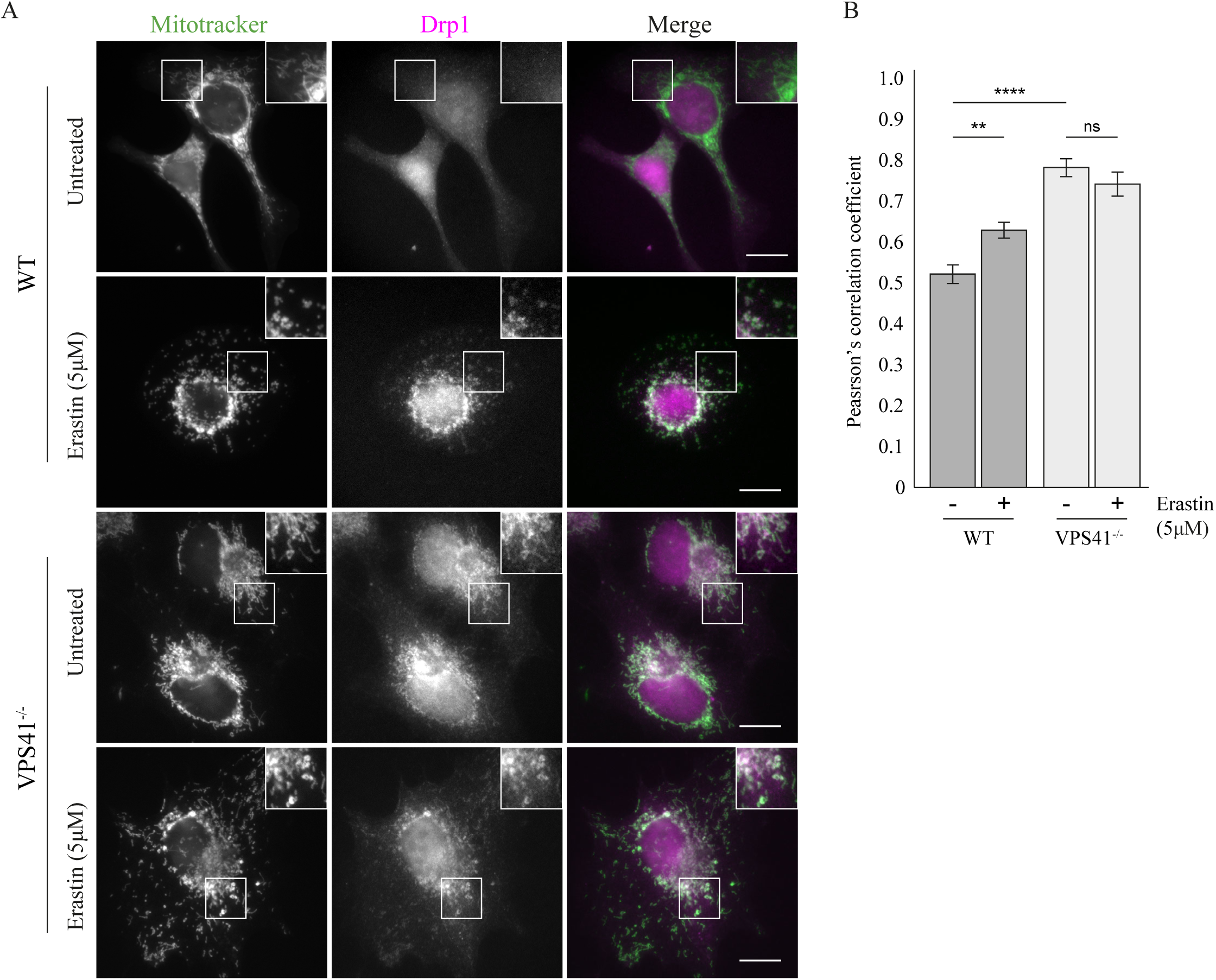
Mitochondrial fission induced by AMPK is increased in VPS41^-/-^ cells. A. Immunofluorescence of endogenous Drp1 in cells incubated with MitoTracker to visualize mitochondria in WT and VPS41^-/-^ cells under basal (Untreated) and Ferroptosis induced (Erastin) conditions. Co-localization between Drp1 and MitoTracker is observed under basal conditions in VPS41^-/-^ cells, whereas WT cells only show a cytosolic distribution of Drp1. Upon Erastin treatment co-localization between Drp1 and MitoTracker is also observed in WT cells. (Scale bar; 10µm) (n=2). B. Quantification of A, co-localization between Drp1 and MitoTracker is significantly higher in VPS41^-/-^cells under basal conditions compared to WT cells. Upon Erastin treatment, co-localization significantly increases in WT cells, whereas no effect is observed in VPS41^-/-^ cells. Bars represent mean ± SEM, **P < 0.01, ****P < 10^-5^, Unpaired t-test.

Taken together, these data showed that Drp1-mediated mitochondrial fission was increased in VPS41^-/-^cells, which explained the severe mitochondrial fragmentation phenotype.

## Discussion

Ferroptosis is a recently discovered type of regulated cell death characterized by intracellular iron overload and ROS accumulation and caused by iron-dependent lipid peroxidation. Ferroptosis has been implicated in e.g. Alzheimer’s disease, Parkinson’s disease and Neurodegeneration with Brain Iron Accumulation (NBIA) disorders(Guiney et al., 2017; Ji et al., 2022; X. Jiang et al., 2021). Recently, a genome-wide CRISPRi/a screen in human neurons linked Ferroptosis to lysosomal failure and identified human VPS41 as risk factor for the onset of Ferroptosis(Tian et al., 2021). Moreover, we and others have found that patients bearing compound heterozygote mutations in VPS41 show iron accumulations in the basal ganglia, globus pallidus and substantia nigra, accompanied by severe neurodegeneration(Sanderson et al., 2021; Steel et al., 2020; Van der Welle et al., 2021). However, how VPS41 depletion affects iron metabolism has never been addressed. As a first step towards understanding the molecular mechanisms underlying this phenomenon, we performed a set of assays to study iron metabolism in VPS41^-/-^ cells of human origin. We showed that absence of VPS41 resulted in the accumulation of a highly reactive, labile Fe^2+^ pool in both the cytosol and aberrant endo-lysosomes. In addition, we showed that VPS41 deletion led to an excess of ROS formation and a severe fragmentation of the mitochondrial network. These findings strengthen the proposed role for VPS41 in iron homeostasis and in addition reveal a severe effect of VPS41 deletion on mitochondrial fission. The data also imply that the neurodegeneration seen in VPS41 patients could at least partially be due to Ferroptosis.

To assess the effect of VPS41 depletion on intracellular iron homeostasis, we analyzed the intracellular protein levels of Transferrin Receptor (TfR), Transferrin (Tf), Ferritin and Ferroportin in Wildtype (WT) versus VPS41 knockout (KO, VPS41^-/-^) A549 cells. Tf and TfR are responsible for uptake of iron from the extracellular space and their protein levels decrease upon intracellular iron overload. Ferritin and the iron exporter Ferroportin control the intracellular labile iron pool of reactive Fe^2+^, and their protein levels increase upon iron overload(Kakhlon & Cabantchik, 2002; Kidane et al., 2006; Lv & Shang, 2018). Interestingly, VPS41^-/-^ cells showed a significant decrease in TfR proteins and a concomitant reduction in Tf uptake. Concurrently, they exhibited significantly higher levels of Ferritin and Ferroportin proteins, all indicators for iron overload. Increased iron levels are known to inhibit IRP/IRE binding to TfR-mRNA, resulting in mRNA degradation and, consequently, reduced TfR protein levels.(Wilkinson & Pantopoulos, 2014) Concomitantly, we found that reducing iron levels in VPS41⁻/⁻ cells through iron chelation led to increased TfR expression and decreased levels of ferritin and ferroportin. This result indicated that the altered expressions of iron-regulatory proteins in VPS41-deficient cells was indeed a consequence of elevated intracellular iron levels and furthermore demonstrated that the IRP/IRE system remains functionally intact in the absence of VPS41.

In agreement with the observed changes in iron regulatory proteins, we also found an increase in the labile iron pool of VPS41^-/-^ cells. Increased levels of the fluorescent, membrane permeable FeRhoNox-1 probe - detecting Fe^2+^ - was found in the cytoplasm, as well as in enlarged endosomes (HOPS bodies) that are induced by the absence of the HOPS complex (van der J. Beek et al., 2022). An increase in Fe^2+^ levels is commonly associated with oxidative stress, since Fe^2+^ catalyzes the formation of toxic reactive oxygen species (ROS) via the Fenton reaction. Excess levels of ROS cause damage to proteins and organelles ultimately resulting in cell death(Nakamura et al., 2019; Rynkowska et al., 2020). Moreover, increased ROS levels induce Drp1-mediated mitochondrial fission and induce the peroxidation of phospholipids containing polyunsaturated fatty acids(Rynkowska et al., 2020; Sun et al., 2018). Using multiple assays, we demonstrated that VPS41 depletion indeed led to increased ROS levels and induced lipid peroxidation. First, we found that the signal emitted by the cellROX probe, indicating ROS, was manifold stronger in VPS41^-/-^ than in WT cells (Fig 3B). Interestingly, the ROS signal appeared as clusters, suggesting accumulation in specific regions of the cells. These could be cytosolic aggregates but may also represent the HOPS bodies(J. van der Beek et al., 2024). In addition, we found that VPS41^-/-^ cells showed a clear increase in mitochondrial Drp1 and concomitantly significantly smaller mitochondria, which is a known effect of increased ROS production(Neitemeier et al., 2017), (M. Gao et al., 2019). Lastly, we measured Malondialdehyde (MDA) levels, a common product of lipid peroxidation of polyunsaturated fatty acids(Ayala et al., 2014). This showed that MDA levels were doubled in VPS41^-/-^ compared to WT cells (Fig 3B). An increase in lipid peroxidation is the net result of glutathione (GSH) depletion and/or inhibition of Glutathione Peroxidase 4 (GPx4), which together catalyze the reduction of lipid peroxides. GSH is the major scavenger of free radicals to prevent cellular damage by ROS. Diminished GSH levels increase the cells’ vulnerability towards oxidative stress. Notably, we found by untargeted metabolomics in patient derived VPS41^S285P/R662*^ fibroblasts, that levels of a metabolite corresponding in m/z to GSH were strikingly reduced, compared to fibroblasts from a healthy, non-related, individual (data not shown). These latter results should be validated on more fibroblast samples and expression of glutathione biosynthetic genes such as GCL could be assessed to help explain the reduced GSH levels observed in patient-derived fibroblasts. Since a reduction in GSH levels suggest that Ferroptosis suppressing genes might be altered, expression of ferroptosis regulators like GPx4 could be investigated.

Collectively, these independent assays all indicated that VPS41⁻/⁻ cells exhibit oxidative stress, including elevated lipid peroxidation, a key driver of Ferroptosis. To our knowledge, this is the first study to show the impact of VPS41 knockout on iron metabolism in human cell lines. Our studies are in agreement with early studies in the yeasts *Saccharomyces cerevisiae* and *Trypanosoma brucei,* in which VPS41 was originally defined as an essential protein for growth on low-iron medium (Lu et al., 2007; Radisky et al., 1997). Deletion of VPS41 in yeast resulted in defective Fet3p activity, a multicopper oxidase located at the plasma membrane that catalyzes Fe^2+^ to Fe^3+^ (Lu et al., 2007; Radisky et al., 1997). Ceruloplasmin, the mammalian homologue of Fet3p, is a ferroxidase that alleviates Ferroportin in exporting iron from the cell(De Domenico et al., 2007; Musci et al., 2014). Activation and plasma membrane localization of Ceruloplasmin is regulated by Wilson disease protein ATP7B, a copper transporter (Harada et al., 2005; Yanagimoto et al., 2009). In analogy, it would be interesting to determine if ATP7B and ceruloplasmin are also affected by VPS41 depletion.

A likely explanation for the observed phenotypes in VPS41^-/-^ cells are defects in intracellular transport. Notably, we found accumulations of Fe^2+^ and possible ROS in HOPS bodies^48^. HOPS bodies are non-acidified, enzymatically inactive, hybrid early-late endosomes that fail to fuse with lysosomes and accumulate a.o. cholesterol(Ndoj et al., 2025) and autophagic substrates(J. van der Beek et al., 2024). Fe^2+^ could be aberrantly maintained inside these altered endosomes, perhaps due to intraluminal lipid and protein accumulations. Alternatively, HOPS bodies might contain reduced levels of iron channels (i.e. DMT1) or Fe^2+^ could be actively confined within endosomes to prevent utilization in the Fenton reaction, which mainly occurs in the cytosol and mitochondria(Sun et al., 2018), (Fernández et al., 2016), (Blaby-Haas & Merchant, 2014). Additionally, we and others have shown that depletion of VPS41 or other HOPS subunits impairs recycling, from HOPS bodies, for example of the mannose 6-phosphate receptor(J. van der Beek et al., 2024). VPS41 knockdown cells(Pols, Ten Brink, et al., 2013), as well as preliminary data from VPS41 knockout cells (J. van der Beek, personal communication), do not show a clear change in TfR localization, but we cannot exclude a potential effect on recycling of TfR or other proteins involved in iron metabolism. Another possible transport defect contributing to the phenotypes described here is reduced autophagic flux, which is known to occur in the absence of a functional HOPS complex due to impaired transport of endocytosed or autophagocytosed material to lysosomes(Van der Welle et al., 2021). Damaged mitochondria are typically degraded by mitophagy, a specialized type of macro-autophagy to remove old or damaged mitochondria. Inefficient degradation of damaged mitochondria might explain the ROS accumulation in VPS41 depleted or mutated cells. Given the striking mitochondrial fragmentation observed, further characterization of mitochondrial function, such as measuring membrane potential and respiration or assembly of the complexes of the electron transport chain would reveal whether VPS41 loss causes broader mitochondrial dysfunction beyond morphology.

Finally, VPS41 is essential for recruiting TGN-derived transport vesicles carrying lysosomal membrane proteins, such as LAMP1 and NPC1. This process involves binding of VPS41 to vesicle-associated GTPase Arl8b, followed by HOPS-dependent fusion with endosomes 33,124,146,339,342. It would be of interest to determine whether the transport of iron- and copper-regulating membrane proteins also depends on this VPS41-dependent biosynthetic pathway.

Alternatively, the role of VPS41 in iron metabolism may be linked to the non-canonical mTORC1 signaling pathway. Previously, we showed that depletion of VPS41 or other HOPS components, prevented phosphorylation of TFEB/TFE3, two members of the MiTF/TFE family of transcription factors. This resulted in their constitutive nuclear localization and consequent activation of the CLEAR network of lysosomal and autophagy genes, whereas phosphorylation of canonical mTORC1 substrates was not affected(Van der Welle et al., 2021). Activation of TFEB/TFE3 and the resulting increase in autophagy has a neuroprotective role in Parkinson’s disease by promoting clearance of misfolded proteins and preventing Ferroptosis by reducing cellular iron overload(Chen et al., 2023). Thus, constitutive activation of TFEB in VPS41 depleted cells ((Van der Welle et al., 2021) and chapter 3 of this thesis) might partially suppress the increase in cellular iron levels caused by VPS41 depletion. Interestingly, depletion of folliculin (FLCN), the causative gene for Birt-Hogg-Dubé (BHD) syndrome, results in a TFEB phenotype similar to that observed after VPS41 depletion(Hong et al., 2010; Lawrence et al., 2019; Napolitano et al., 2020, 2022). An iron-rich diet reversed the disease phenotype in a *Drosophila* model of BHD, suggesting a role for FLCN in iron uptake(Wang et al., 2021). This finding is particularly significant, since we recently discovered that VPS41 and other HOPS subunits interact with FLCN (chapter 3 of this thesis). These interactions are required for lysosomal localization of FLCN, which is necessary for mTORC1 dependent phosphorylation of TFEB/TFE3 to prevent transport to the nucleus. It is therefore tempting to speculate that the role of VPS41 in iron homeostasis might require interactions with FLCN.

FLCN is also an interactor of AMPK, the central regulator of energy homeostasis(Ramirez Reyes et al., 2021). AMPK is activated by mitochondrial stress resulting in phosphorylation of Mitochondrial Fission Factor (MFF), which enables recruitment of Drp1 to catalyze mitochondrial fission(F. Gao et al., 2021; Toyama et al., 2016). AMPK phosphorylates TFEB independent of mTORC1, on different serine residues, to enable transcriptional activity345,346. Whether AMPK is involved in the proposed VPS41-FLCN axis for iron homeostasis is an interesting question for future research. Also, it remains to be established what the order of events is; does iron overload result in ROS production and enhanced mitochondrial fission, or is the sequence of events different?

Iron homeostasis is especially important for neuronal development, neurotransmitter synthesis, synapse establishment and myelin formation347. Neurons are therefore extremely vulnerable to ROS and many forms of neurodegenerative diseases are associated with oxidative stress140,317,348. VPS41 was identified as neuroprotector in Alzheimer’s and Parkinson’s disease, due to its role in transporting endocytosed and autophagy cargo’s to degradative lysosomes(Griffin et al., 2018; Harrington et al., 2012; Ruan et al., 2010). This study indicates an additional role for VPS41 in iron homeostasis, ROS production and mitochondrial morphology, providing an interesting basis for future research, especially in neuronal cells.

## Methods and Materials

### Cell culture and light microscopy

The generation of the CRISPR/Cas9 knockout cells is described in (Van der Welle et al., 2021). Hela WT and VPS41^-/-^ cells were cultured in Dulbecco’s Modified Eagle’s Medium (Invitrogen), supplemented with 10% FCS and 1% Penicillin/streptomycin (Invitrogen) at 37°C, 5% CO2. A549 WT and A549 VPS41^-/-^ cells were cultured in RPMI-1640 medium (ThermoFisher) supplemented with 10% FBS, 1% Penicillin/streptomycin and 1% L-Glutamine (ThermoFisher). Iron chelation was performed by overnight incubation (16 hours) with 100µM 2,2’-Bipyridyl (Sigma) added to cell culture medium. Ferroptosis was induced by overnight treatment with 5µM Erastin (Sigma), added to culture medium.

Transferrin (Tf) uptake was assessed using cells cultured on coverslips in a 24-well plate and incubated with Tf conjugated to Alexa-488 (Life Technologies) for 2, 10 or 20 minutes at 37°C. Cell were fixed with 4% PFA in PBS at 4°C for 1 hour, placed on glass slides with 5 µL Prolong gold DAPI (Life Technologies), cured overnight at room temperature and imaged using the DeltaVision (GE Healthcare Life Sciences). Quantification of Tf uptake was quantified using Volocity software.

### FeRhoNox-1

A549 Cells cultured on coverslips in a 24-well plate were incubated with 5 µM FeRhoNox-1 (tebu-bio, GC901) in Hanks’ Balanced Salt Solution (HBSS, Sigma) for 60 minutes at 37 °C. The cells were washed 3 times with HBSS and fixed for 15 minutes with 4% w/v paraformaldehyde (PFA, Polysciences Inc.). After fixation, cells were washed 3 times with PBS prior to immunofluorescence labeling.

### CellROX Deep Red

A549 cells cultured in ibidi µ-Slide 8 Well Chambers were incubated with 5µM cellROX Deep Red Reagent (ThermoFisher, C10422) added to the cell culture medium for 30 minutes at 37 °C. Cells were washed 3 times with culture medium and imaged at the DeltaVision (GE Healthcare) at 37 °C and 5% CO2.

### MitoTracker Red CMXRos

Cells cultured on coverslips in a 24-well plate were incubated with 100nM MitoTracker Red for 15 minutes at 37 °C, washed 3 times with culture medium and then fixed for 15 minutes with 4% PFA in PBS. After fixation, cells were washed 3 times with PBS prior to immunofluorescence labeling. Quantification of mitochondrial branch length was performing using the MiNA (Mitochondrial Network Analysis) ImageJ macro. For colocalization analysis Volocity software was used.

### Immunofluorescence labeling

Cells cultured on coverslips were fixed using 4% PFA in PBS, washed 3 times with PBS and permeabilized with 0,1% Triton X-100 (Sigma) in PBS for 10 min. Cells were blocked with 1% BSA (Roche) in PBS for 15 minutes. The coverslips were incubated for 1 hour with primary antibodies diluted in 1% BSA in PBS, washed 3 times in PBS and incubated with secondary antibody, diluted in 1% BSA in PBS, for 1 hour. Coverslips were washed 3 times with PBS and placed on slides with Prolong gold DAPI (Life Technologies) and cured overnight. Antibodies and their specific dilutions used in this study are specified in Table 1.

**Table 1.** Antibodies and dilutions used.

| Primary Antibody | Use | Dilution | Company | Catalog Number |
| --- | --- | --- | --- | --- |
| M $\alpha$ TfR | WB | 1:1000 | ThermoFisher | 13-6890 |
| R $\alpha$ Ferritin | WB | 1:1000 | Abcam | 75973 |
| R $\alpha$ Ferroportin | WB | 1:1000 | Invitrogen | PA5-22993 |
| M $\alpha$ Actin | WB | 1:2000 | MP Biomedicals | 69100 |
| R $\alpha$ FLCN | WB | 1:4000 | Cell Signaling | 3697 |
| R $\alpha$ AMPKα | WB | 1:1000 | Cell Signaling | 2532 |
| R $\alpha$ phospho-AMPKα | WB | 1:1000 | Cell Signaling | 2535 |
| G $\alpha$ Cathepsin D | IF | 1:500 | R&D Systems | AF1014 |
| R $\alpha$ EEA1 | IF | 1:150 | Cell Signaling | C45B10 |
| R $\alpha$ Rab7 | IF | 1:200 | Cell Signaling | 9367 |
| M $\alpha$ Tomm20 | IF | 1:750 | BD Transduction Laboratories | 612278 |
| M $\alpha$ Drp1 | IF | 1:250 | Santa Cruz Biotechnology | sc-271583 |
| Secondary Antibody | Use | Dilution | Company | Catalog Number |
| IRDye® 800CW G $\alpha$ M - IgG | WB | 1:5000 | LI-COR | 925-32210 |
| IRDye® 800CW G $\alpha$ R - IgG | WB | 1:5000 | LI-COR | 925-32211 |
| IRDye® 680CW G $\alpha$ R - IgG | WB | 1:5000 | LI-COR | 925-68071 |
| D $\alpha$ G-ALEXA488 | IF | 1:250 | ThermoFisher | A-11055 |
| D $\alpha$ R-ALEXA488 | IF | 1:250 | ThermoFisher | A-21206 |
| R $\alpha$ M-ALEXA488 | IF | 1:250 | ThermoFisher | A-11061 |
| G $\alpha$ M-ALEXA488 | IF | 1:250 | ThermoFisher | A-11029 |

### Lipid peroxidation (MDA) assay

Lipid peroxidation was determined by assessing the malondialdehyde (MDA) levels using the Lipid Peroxidation (MDA) Assay Kit (Colorimetric/Fluorometric) (Abcam) according to the manufacturer’s protocol. Briefly, cells were harvested, lysed and homogenized using a Dounce homogenizer. Samples were incubated with Thiobarbituric Acid for 1 hour at 95°C and added to a 96-well microplate. Absorbance was measured at OD 532nm.

### Western blotting

Cells were grown in a 6 cm petri dish till 90% confluence was reached, placed on ice, washed with cold PBS and lysed using lysisbuffer (1M Tris pH 7.5, 4M NaCl, 0.5M EDTA, 1M NaF and 10% Triton X-100) complemented with protease inhibitor and 2M DTT. Cells were scraped and protein levels were equalized using a Bradford Protein Assay (BioRad). Samples were run on 4-15% pre-cast gels (BioRad) and transferred using Trans-Blot® Turbo™ RTA Mini PVDF Transfer Kit and the Trans-Blot® Turbo™ Transfer system (Bio-Rad). Membranes were blocked using Odyssey® Blocking Buffer (LI-COR) in 0.1% TBST for 1 hour prior to incubation with primary antibody diluted in Odyssey® Blocking Buffer in 0.1% TBST overnight at 4°C. Membranes were washed extensively using 0.1% TBST and incubated with secondary antibody, also diluted in Odyssey® Blocking Buffer in 0.1% TBST, for 1 hour at room temperature. Membranes were again washed extensively with 0.1% TBST, followed by 2 washing steps in PBS prior to scanning the membranes using the Amersham Typhoon 5 (GE healthcare). Antibodies and their specific dilutions used in this study are specified in Table 1.

### Transmission electron microscopy (TEM)

Cells were fixed with a combination of 2% paraformaldehyde and 2.5 % glutaraldehyde overnight at 4°C. After fixation, cells were washed with 0.1M PB 3 times for 5 minutes and supplemented with 1 mL 0.1M PB with 2% gelatin. The cells were scraped and spun down in at 9000 rpm for 1 minute. The supernatant was removed and cells were resuspended in 2% low melting point agarose and spun down for 1 minute at 9000 rpm. The agarose was cut and transferred to a 5 mL glass vial with 0.1M PB. Post fixation was performed with 1% OsO4 and 1.5% K3Fe (III) (CN)6 in 0.1M PB for 1 hour at 4°C and 0.5% uranyl acetate in dH2O for 1 hour at 4°C in the dark. Samples were passed through a graded acetone : dH2O series for dehydration (70% overnight at 4°C, 90%, 96%, 1×15 minutes each, and 100% 3×30 minutes each). Epon resin infiltration was done (25%, 50%, 75% in acetone for 45 minutes each), followed by 2×45 minutes infiltration steps using 100% Epon resin. Final embedding was performed in 100% Epon resin for 72 hours at 60°C. Sectioning was performed using a Leica UCT/FCS and imaging was performed on a FEI Tecnai 12 80 kV transmission electron microscope. Stitch reconstruction was performed with the Etomo software. Quantifications on mitochondrial area was performed using ImageJ.

### Mitochondrial 3D reconstruction

Mitochondria from Hela WT and VPS41^-/-^ cells were reconstructed using a Focused Ion Beam Scanning Electron Microscope (FIB-SEM) dataset described in (Fermie et al., 2018). Reconstruction was performed with 3dmod.

### Metabolomics

Untargeted metabolomics analysis of fibroblasts was performed using direct infusion high-resolution mass spectrometry according to Haijes et al. Each sample was measured in triplicate by injecting from the same well. Data processing was performed by an in-house metabolomics pipeline in R (Source code available at: https://github.com/UMCUGenetics/DIMS). Mass peaks were annotated by matching the mass over charge ratio (m/z) to metabolite masses in the Human Metabolite Database (HMDB) within a range of 5 parts per million. For each sample, the mean intensity of the triplicate injection was reported for each mass peak.

### Statistics

Quantification of all western blots was performed using Fiji software (Schindelin et al., 2012). Two-tailed Students t tests were performed to compare two samples whereas ANOVA was performed for comparison of multiple samples to analyze statistical significance. Data distribution was assumed to be normal, but this was not formally tested. All error bars represent the Standard Error of the Mean (SEM) as indicated in the specific figure legends.

## Appendix

We thank our colleagues of the Cell Biology section and Center for Molecular Medicine for the fruitful discussions and feedback. We especially thank Dr. Roberto Zoncu for this insightful comments and valuable input.

Author contribution: Experiments: RW, RJ, JJ; Study design: RW, RJ, JJ, NV and JK; Manuscript reviewing and revision: RW, JJ, NV; Study design and supervision: RW, JJ, NV and JK; Funding: JK; Manuscript writing: RW and JK.

The authors declare no conflict of interest.

